# Intra and inter-annual climatic conditions have stronger effect than grazing intensity on root growth of permanent grasslands

**DOI:** 10.1101/2020.08.23.263137

**Authors:** Catherine Picon-Cochard, Nathalie Vassal, Raphaël Martin, Damien Herfurth, Priscilla Note, Frédérique Louault

**Author notes:** **Cite as:** Picon-Cochard C, Vassal N, Martin R, Herfurth D, Note P, Louault F (2021) Intra and inter-annual climatic conditions have stronger effect than grazing intensity on root growth of permanent grasslands. bioRxiv, 2020.08.23.263137, version 6 peer-reviewed and recommended by PCI Ecology. https://doi.org/10.1101/2020.08.23.263137. **Posted:** MS ID#: BIORXIV/2020/263137 16 03 2021. **Recommender:** Jennifer Krumins. **Reviewers:** Three anonymous reviewers. This article has been peer-reviewed and recommended by *Peer Community in Ecology* https://doi.org/10.24072/pci.ecology.100073.

## Abstract

**Background and Aims:** Understanding how direct and indirect changes in climatic conditions, management, and species composition affect root production and root traits is of prime importance for the delivery of carbon sequestration services of grasslands. This study considers the effects of climatic variability and gradients of herbage utilisation by grazing on root production over the course of two years. The root and leaf traits of the plant communities were determined to detect their capacity to predict above- and below-ground net primary production, ANPP and BNPP, respectively.

**Methods:** A long-term field experiment was used to compare the effects of abandonment and low (Ca-) and high (Ca+) grazing intensities (resulting in mean residual plant heights of 15.2 cm and 7.7 cm, respectively) induced by grazing rotations on upland fertile grasslands after 10 years of treatment application. Ingrowth cores and exclusion cages were used to measure, respectively, the root and shoot mass production several times each year and at an annual scale. The root and leaf traits of the communities were measured near the vegetation’s peak growing season.

**Results:** We observed strong seasonal root production across treatments in both a wet and a dry year, but the response to grazing intensity was hardly observable within growing seasons. In the abandonment treatment, the spring and autumn root growth peaks were delayed by approximately one month compared to the two cattle treatments, possibly due to a late plant canopy green-up induced by lower soil temperatures and an accumulation of litter. The BNPP was slightly lower in the abandonment treatment compared to the cattle treatments only during the dry year, whereas a decline of the ANPP in the abandonment treatment compared to the Ca+ treatment was observed during the wet year. In response to drought, which occurred during the second year, the root-to-shoot biomass ratio was stable in the cattle treatments but declined in the abandonment treatment. The higher allocation to root mass could benefit plant communities under drier conditions.

**Conclusions:** Rotational grazing pressures and climatic condition variabilities had limited effects on root growth seasonality, although drought had stronger effects on the BNPP than on the ANPP. The stability of the root-to-shoot biomass ratio during the dry year evidenced a higher resistance to drought by grazed versus abandoned grassland communities.

## Introduction

Permanent grasslands provide many services that connect to human activities through livestock products but also contribute to regulate greenhouse gas emissions because their soils accumulate large amounts of carbon (C) in organic matter fractions. Root activity (growth, exudation, and turnover) contributes to C and nitrogen (N) inputs and to absorb both nutrients and water, which are essential to fix atmospheric carbon dioxide (CO2) and produce plant biomass. The intensification of management practices may affect these services as well as climate variability (Conant et al., 2001; Jones and Donnelly, 2004; Soussana and Duru, 2007). Thus, improving our understanding of grassland roots dynamics under different management and climatic conditions may help to identify management options to maintain the forage production and C sequestration abilities of this ecosystem and thus its sustainability.

Different management practices modify forage production and the amount of soil C and N through both the direct effects of defoliation, fertilisation, or returns of excreta to soil on root growth and soil abiotic factors and the indirect effects of species composition changes (Bardgett and Wardle, 2003; Dawson et al., 2000; Soussana et al., 2004). In mown grasslands, the root mass production is generally lower when grasses are frequently mown and fertilised (Leuschner et al., 2013; Picon-Cochard et al., 2009). This result may be explained by changes in the root-to-shoot allocation with an increase of above-ground growth to maximise light capture. The complexity of these phenomena in grazed grasslands is greater than in mown systems owing to animals’ selective defoliation of plant species and because returns to soil are spatially heterogeneous (Rossignol et al., 2011). In addition, the level of soil fertility may buffer the degree of root response to defoliation in grazed grasslands as plants exhibit specific responses to defoliation in fertile and unfertile grasslands (Duru et al., 1998). Overall, the complexity of phenomena being direct and indirect effects on plants could explain why no clear trend has been found for the effects of grazing on above- and below-ground production (e.g., refer to the syntheses of Milchunas and Lauenroth [1993] and McSherry and Ritchie [2013]), although two meta-analyses have emphasised the negative effect of grazing intensity on above- and belowground C stocks compared to ungrazed systems (Zhou et al., 2017; Li et al., 2018). Furthermore, repeated defoliations induced by grazing or mowing grasslands can simultaneously increase soil temperatures and moisture (Moretto et al., 2001; Pineiro et al., 2010; Smith et al., 2014). Soil moisture can also be modified by high stocking rates through changes in the soil bulk density due to soil compaction and changes in the leaf area index after defoliation (Pineiro et al., 2010). These direct effects of grazing on soil abiotic factors should affect the root growth of grazed grasslands, although these phenomena have not been well documented in field conditions.

Species composition change induced by management is also an important determinant of above- and below-ground responses in grazed grasslands. Intensive practices (e.g., high grazing intensity or fertilisation) generally favour the development of fast-growing species (plant exploitative strategy), whereas the opposite extensive practices (e.g., low grazing intensity or the absence of fertilisation) favour slow-growing species (plant conservative strategy; Klumpp et al., 2009; Louault et al., 2005; Soussana and Lemaire, 2014; Wardle et al., 2004). The root-to-shoot biomass allocation, as well as the functional traits (used as proxies of ecosystem properties such as the ANPP or BNPP; e.g., Laliberté and Tylianakis, 2012), are thus likely to change in response to an intensification of practices, such as from ungrazed to intensively grazed temperate grasslands, alpine meadows, steppes or desert steppes (Klumpp and Soussana, 2009; Zeng et al., 2015). According to Ziter and MacDougall (2013), the uncertainty surrounding nutrient-defoliation responses makes it difficult to predict whether C storage would be higher in managed versus unmanaged grasslands. Thus, soil fertility should be considered when comparing different grazing intensities in grasslands (Louault et al., 2005).

Increased climate variability is another source of response uncertainty in managed ecosystems. As more frequent and longer periods of drought associated with heat waves may threaten and shape the long-term dynamics of perennial ecosystems such as grasslands (Brookshire and Weaver, 2015), it is important to understand how above- and below-ground compartments respond to climate variability. However, the data on above- and below-ground biomass responses to drought for grasslands are limited (Byrne et al., 2013; Wilcox et al., 2015; Li et al., 2018), although some evidence shows that resource conservation strategy is associated with drought tolerance (Pérez-Ramos et al., 2012; Reich, 2014). Changes in root morphology and functioning may thus be important determinants in plants’ adaptive strategies to drought and have been less studied than above-ground plant responses (Biswell and Weaver, 1933; Dawson et al., 2000; McInenly et al., 2010). However, there are not enough data to make generalisations about the combined impacts of management and climatic condition variabilities, such as precipitation reduction, on root and shoot biomass production and plant traits defining plant strategies related to resource use and grazing intensity.

This study comprised of a long-term field experiment for which controlled grazing intensity was applied for 10 years. We compared grazing abandonment and two levels of herbage utilisation by grazing based on five rotations per year. In two consecutive years, the ingrowth core method was used to measure the monthly root biomass production and calculate the BNPP. The ANPP was measured using grazing exclusion cages, and the community-weighted mean (CWM) leaf and root traits were assessed the first year. We tested the following hypotheses: (i) high grazing intensity increases above-ground mass at the expense of root production as a result of the direct negative effect of defoliation on root growth, whatever the climatic conditions; (ii) interannual climatic conditions modulate the above- and below-ground biomass production response to grazing intensity as a consequence of the higher presence of defoliation-tolerant and drought-sensitive species (*Lolium perenne* or *Trifolium repens*) in the high grazing intensity treatment; (iii) root traits respond to treatments and are a determinant of the BNPP, as observed for leaf traits and the ANPP.

## Methods

### Site characteristics

The experiment transpired at the long-term observatory network (ACBB, https://www.anaee-france.fr/en/infrastructure-services/in-natura-experimentation/agrosystem/acbb) located in St-Genès-Champanelle, France (45° 43’N, 03° 01’E, 880 m a.s.l.). The local climate is semi-continental with oceanic influences and a mean annual temperature and precipitation of 8.5°C and 784 mm, respectively (Table 1). The site supports mesotrophic multi-specific permanent grasslands dominated by species with high Ellenberg indicator values for N (Schaffers and Sykora, 2000), indicating a high level of fertility for the site (Table S1; Louault et al., 2017). The soil is a cambisol with a sandy loam texture developed on a granitic bedrock. Differences in the local soil composition and profile led us to consider two distinct blocks characterised by an eutric cambisol (54% sand, 26% silt, 20% clay, 7.0% organic matter, and pH: 5.9) and a colluvic cambisol (50% sand, 26% silt, 24% clay, 7.4% organic matter, and pH: 6.0), including some volcanic materials.

**Table 1.** Air temperature (°C), precipitation (P, mm), potential evapotranspiration (PET) and climatic water balance: cumulated (P – PET, mm) and calculated for the 28 y period 1986-2013, mean values ± SD) and measured for the 10 dates in 2014 and 2015 corresponding to measurements of root growth and averaged (temperature) or summed (P, PET, P – PET) at annual scale.

| Year | Dates | Air temperature | Precipitation | PET | P-PET |
| --- | --- | --- | --- | --- | --- |
|  | **Annual long-term average** | **$8.5 \pm 0.6$** | **$784 \pm 1376$** | **$693 \pm 96$** | **$91 \pm 195$** |
| **2014** | December 12 – February 23 | 3.7 | 98 | 37.5 | 60.5 |
|  | February 24 – March 23 | 5.3 | 27 | 46.3 | -19.3 |
|  | March 24 – April 21 | 7.2 | 23.5 | 68.7 | -45.2 |
|  | April 22 – May 25 | 9.2 | 79.5 | 103.1 | -23.6 |
|  | May 26 – June 22 | 14.2 | 58 | 110.2 | -52.2 |
|  | June 23 – July 20 | 15.1 | 136.5 | 93.9 | 42.6 |
|  | July 21 – August 24 | 14.4 | 90.5 | 100.5 | -10 |
|  | August 25 – September 29 | 13.7 | 141.8 | 79.5 | 62.3 |
|  | September 30 – October 29 | 11.7 | 69 | 36.3 | 32.7 |
|  | October 30 – December 14 | 5.3 | 111 | 10.9 | 72.1 |
|  | **Annual** | **9.2** | **876** | **691** | **157.7** |
| **2015** | December 15 – March 1 | 1.3 | 132.5 | 31 | 101.5 |
|  | March 2 – March 29 | 4.5 | 36.5 | 36.8 | -0.3 |
|  | March 30 – April 23 | 8.5 | 17.5 | 66.4 | -48.9 |
|  | April 24 – May 28 | 11.0 | 66 | 113.6 | -47.6 |
|  | May 29 – June 28 | 15.5 | 62.5 | 129.1 | -66.6 |
|  | June 29 – July 23 | 21.1 | 26 | 136 | -110 |
|  | July 24 – August 27 | 16.4 | 94.5 | 124.6 | -30.1 |
|  | August 28 – September 24 | 12.8 | 77 | 66.3 | 10.7 |
|  | September 25 – October 29 | 7.8 | 55 | 36.1 | 18.9 |
|  | October 30 – December 11 | 7.0 | 54.5 | 25.1 | 29.4 |
|  | **Annual** | **9.4** | **585** | **766** | **-180.9** |

### Management

Prior to the installation of this experiment in 2005, the study area had been used for intensive hay and silage production (combining grazing, mowing, and fertilisation) with mineral fertilisation, and two years preceding the start of the experiment (2003 and 2004), the grassland site was mown three times per year without fertilisation. From 2005, the grassland was managed for 10 years with a gradient of grazing intensity resulting from three treatments: abandonment, low herbage utilisation (Ca-), and high herbage utilisation (Ca+). The Ca- and Ca+ treatments were obtained by modifying the stocking density (6.9 and 13.8 livestock units (LSU) ha^−1^, respectively) with five grazing rotations each year occurring at mid-April, late May, early July, September, and November and lasting on average 9.6, 9.0, 10.7, 8.6, and 2.1 days, respectively. The two cattle treatments corresponded to two levels of herbage utilisation by grazing and had on average a residual plant height of 15.2 cm ± 0.5 (mean ± SE) for Ca- and 7.7 cm ± 0.2 for Ca+ at the end of each grazing rotation. For each treatment, two replicate plots were created per block, resulting in four replicates per treatment and a total of 12 plots (2 blocks x 2 plots x 3 treatments). The average distance between the two blocks was approximately 230 m, and all the treatments were randomised within each block. The size of the plots differed according to the treatment: 2,200 m^2^ for the two cattle treatments and 400 m^2^ for the abandonment treatment.

### Climatic and edaphic conditions

The daily precipitation (mm) and air temperature (°C) were measured for the two years and recorded with an onsite meteorological station. A climatic water balance was calculated as precipitation minus the potential evapotranspiration (P – PET, mm) with the Penman-Monteith equation. The daily soil temperature (°C) was measured with thermocouple sensors (homemade copper-constantan sensors) inserted at a 20-cm depth in each plot and recorded with an HOBO data logger (U12-014, Onset Instruments, MA, USA). The daily volumetric soil water content (SWC; m^3^ m^−3^) of each plot was measured with two probes (ECHO-10, Decagon, USA) inserted horizontally at a 20-cm depth and connected to data loggers (EM5 and EM50, Decagon, USA). From January 2014 to November 2015 (DOY 132–326), the SWC was measured every 30 min and averaged at a daily scale. For each plot, the average values of the two probes were used. The daily relative soil water content (RSWC) data are shown and calculated as the ratio: 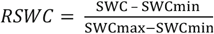, where SWC is the soil moisture at a given day, SWCmin is the minimum value of soil moisture, and SWCmax is the maximum value of soil moisture, all observed during the two years. For the soil temperature and RSWC, the values were averaged according to the root growth timescale.

### Root growth and root mass

Six months before the start of the experiment, shallow (0-20 cm) soil was collected on each of the two blocks, sieved (using a 5 mm mesh size) to remove stones and coarse organic matter, and then left unused outside covered under a shelter and protected from direct sunlight. This air-dried soil was subsequently used to fill the ingrowth cores each month. In December 2013, soil cores were collected with an auger (which had an 8-cm diameter and 0-20 cm depth) for each of the 12 plots at four locations representative of the plant community in the treatment. On average, the mean distance between locations was 19.8 m ± 0.2 for Ca+, 21.7 m ± 0.1 for Ca-, and 17.2 m ± 0.2 for Ab (mean ± standard deviation; refer to Figure S1). After the core harvest, each hole was filled with an 8-mm mesh sized plastic net containing a fixed volume of air-dried sieved soil (called thereafter ingrowth core) collected six months beforehand. For the next two years and approximately each month (2 x 10 times), the ingrowth cores, containing soil and the root and rhizome material that had grown therein, were extracted and then replenished with another fixed volume of air-dried sieved soil. Thus, the monthly and annual root production (BNPP, g m^−2^ y^−1^) were measured from February 2014 to December 2015. The root production period ranged on average 36.5 days, but with longer periods in the winter and shorter periods in the spring and summer (Table 1). In periods without precipitation, a fixed volume of water was added to adjust soil humidity to field conditions. After collection, the ingrowth cores were transported to the laboratory and immediately stored at 4°C before being processed in the next five days. The roots were washed under tap water with a 200-μm sieve and then oven dried for 48 h at 60°C. To measure the root mass stock, the soil cores were collected three times (December 2013 and March and June 2014) with the same auger and near the ingrowth core locations. These samples were stored in the freezer at −18°C; after defrosting, the roots were washed with the same procedure as that used for the ingrowth cores and then oven dried for 48 h at 60°C.

### Root traits

Subsamples of the washed roots collected with the ingrowth cores in June 2014 were fresh weighed and then frozen at −18°C before performing a morphology analysis. After defrosting, the roots were stained with methylene blue (5 g L^−1^) for approximately 5–10 minutes, rinsed in water, spread in a transparent glass box containing a thin layer of water, and covered with a transparent plastic sheet. High resolution images were recorded with a double light scanner (800 dpi, perfection V700, Epson, JA) and analysed using WinRhizo software (PRO 2012b, Regent Instruments, CA) with the automatic procedure. Two scans per location were recorded and separately analysed to measure the root length (m), root volume (cm^3^), root surface area (m^2^), average root diameter (mm), and root length by class diameter (there are 13 classes: 11 with a 0.1-mm interval and 2 with a 0.5-mm interval). The specific root length (SRL; m g^−1^), root tissue density (RTD; g cm^−3^), and specific root area (SRA; m^2^ g^−1^) were calculated for fine roots as in Picon-Cochard et al. (2012).

### Botanical composition

The species contribution (%) was visually observed on a 20-cm diameter circle around each ingrowth core location in April (cattle treatments) and May (abandonment treatment) 2014. For each zone, a score on a 10-point scale was allocated to the species present according to their volume occupancy, and the percentage of each species was calculated at the plot scale by averaging the values of the four zones. Table S2 lists the species and their relative contributions.

### Above-ground biomass production

On each plot and on each sampling date, four fenced sampling areas (0.6 × 0.6 m) were used to measure the accumulation of above-ground biomass after the above-ground standing biomass was clipped to 5.5 cm, oven dried, and weighed. Measurements were made five times over the year, once before each grazing event in the Ca+ and Ca-treatment plots and three times (spring, summer, and autumn) in the abandonment plots. The sampling areas were moved within the plot at each measurement date during the year. The annual aboveground net primary production (ANPP, g m^−2^ y^−1^) was calculated as the sum of the successive biomass accumulations throughout the year.

### Leaf traits

The CWM trait values of the leaf dry matter content (LDMC), specific leaf area (SLA) and reproductive plant height were calculated for each ingrowth core zone using both the relative contribution of the dominant species to the community (i.e., species that account for at least 85% of the cumulated species contribution to the community) measured in 2014 and the leaf trait measurements made at plot scale in 2006 and 2007. The traits were measured on 10 vegetative plants using standard protocols (refer to the methods in Louault et al., 2005). The reproductive plant height was measured on mature plants located in fenced zones to allow full plant development. The CWM is expressed with the following equation: CWM = ∑ *p_i_* × trait_*i*_, where *p_i_* is the relative contribution of species *i* to the community and trait_*i*_ is the trait of species *i*.

### Statistical analyses

For a given date, data averages of the root mass and root traits collected at each of the four locations (four ingrowth cores in each plot) were used to obtain a single value for each of the 12 plots and test for the effect of the treatment and dates. Before variance analysis (ANOVA), the normality of residuals was inspected with quantile-quantile plots of model residuals, and variance homogeneity was confirmed by checking the plots of model residuals versus the fitted values. Data were transformed if they deviated from the ANOVA assumptions (square root, ln, reciprocal). Linear mixed effects models as available in the R ‘nlme’ package (Pinheiro et al., 2015) were used to perform repeated measure ANOVAs to test the effects of treatments and dates, as well as their interactions on values of root growth, soil temperature, RSWC, and root mass stock, with plots nested within block as a random factor accounting for temporal pseudo-replication (S4). For the root growth dynamics, soil temperature, and RSWC (Figure 1; Table S3), the dates correspond to 20 dates, whereas for the root mass stock, the dates correspond to three harvest dates (Table 2). For the BNPP, ANPP, and root-to-shoot ratio (BNPP/ANPP), the data were analysed using a nested mixed model procedure with the treatments and year used as fixed factors and with plot nested within block used as random factors. For the leaf and root traits data, the treatments were used as fixed factors with plots nested within block used as a random factor. Post-hoc tests were performed to compare the significance levels across fixed factors with a Tukey test (‘lsmeans’ package). Principal component analyses (PCAs) were performed for each year to analyse the relationships between the leaf and root traits, soil temperature, RSWC, root mass stock, ANPP, and BNPP measured at the plot level; the treatments were considered supplementary categories (‘FactoMineR’ package). This statistics approach allows sets of trait and property relationships to be compared to detect response and effect traits and to analyse multiple dimensions of trait relationships, which is not possible with pairs of correlations. All statistical analyses were performed in an R environment (Version 3.5.2, R Core Team, 2012) using RStudio (Version 1.1.463). The scripts are shown in S4.

**Figure 1.**
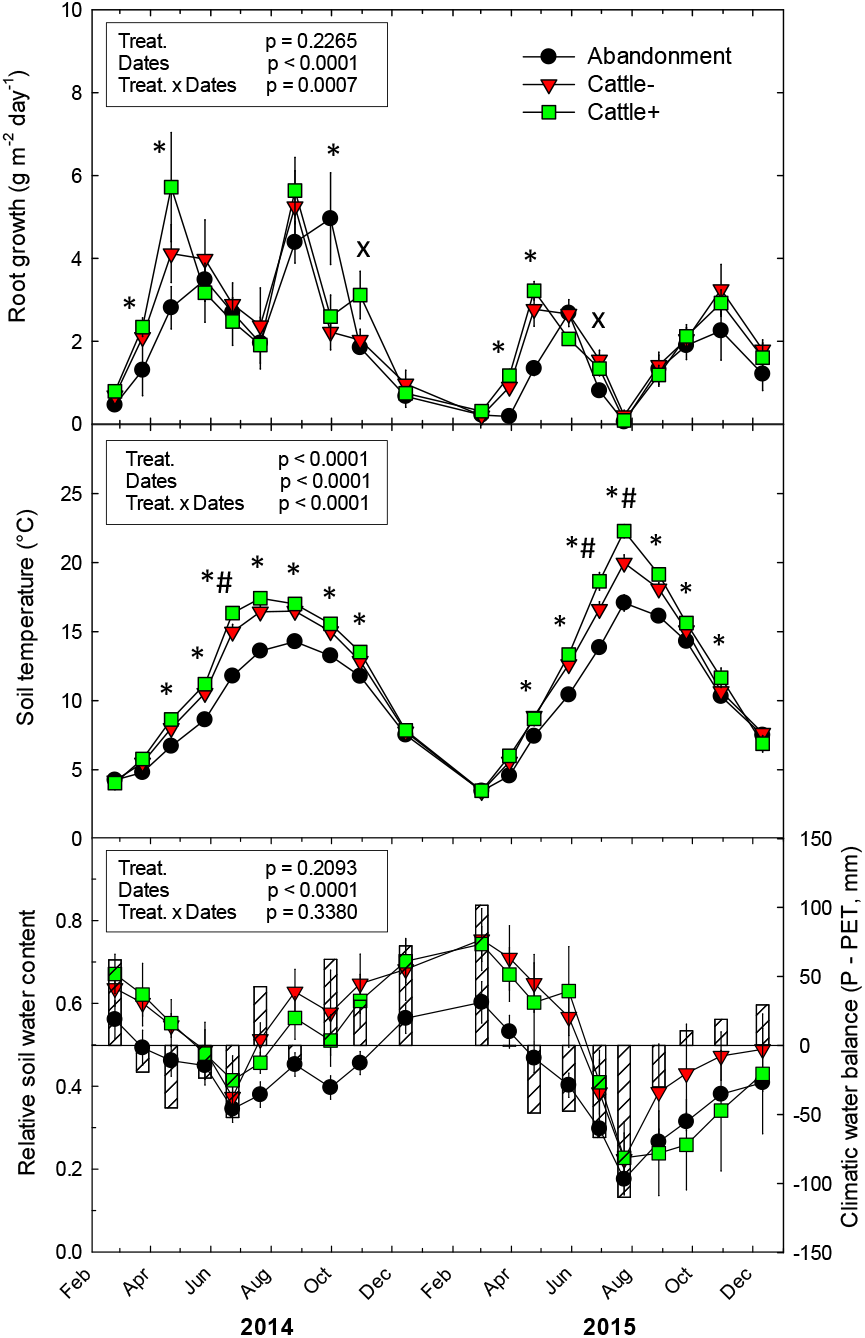
The root growth (g m^−2^ day^−1^), soil temperature (°C), relative soil water content, and climatic water balance (P – PET, mm; hashed bars) dynamics measured over two years for the abandonment, low (Cattle-), and high (Cattle+) grazing intensity treatments. The vertical bars correspond to 1 SE (n = 4). The insets indicate P values from the repeated measure two-tailed ANOVA, where treat. is the treatment; dates, and interaction for the main treatments. *: P < 0.05; x: P ≤ 0.1. For soil temperature, *# corresponds to significant differences between all treatments (Abandonment < Cattle- < Cattle+).

**Table 2.** a) Repeated measure ANOVA is shown for treatment, date (December 2013, March 2014, June 2014) and interaction effects on root mass (g m^−2^). Numerator (num), denominator (den) of degree of freedom (DF) and *F* values are shown. b) Root mass (g m^−2^) of abandonment, low (Cattle-) and high (Cattle+) stocking density treatments measured in winter (December 12 2013), spring (March 20 2014), summer (June 20 2014) and averaged across the three dates. Means ± SE are shown, n = 4. Superscripts ^ns^ correspond to P > 0.05.

| a) | num/den DF | $F$-value |
| --- | --- | --- |
| Treatment | 2/8 | 1.151<sup>ns</sup> |
| Date | 2/18 | 2.027<sup>ns</sup> |
| Treatment × date | 4/18 | 1.340<sup>ns</sup> |

| b) Date | Abandonment | Cattle- | Cattle+ |
| --- | --- | --- | --- |
| December 2013 | 636.4 ± 133.1 | 403.3 ± 66.4 | 496.5 ± 20.6 |
| March 2014 | 559.1 ± 166.2 | 609.2 ± 45.3 | 719.8 ± 47.5 |
| June 2014 | 574.2 ± 84.8 | 482.2 ± 38.6 | 591.2 ± 101.7 |
| 3 dates average | 589.9 ± 99.9 | 498.2 ± 43.6 | 602.5 ± 44.4 |

## Results

### Climatic conditions during the experiment

Compared with the average long-term climatic data for the site, the first and second years of the experiment received higher (+92 mm) and lower (−199 mm) precipitation, respectively (Table 1). The PET in the second year was also higher than the long-term average (a 73-mm difference), leading to a negative annual climatic water balance (P – PET = −181 mm and a deficit of 271 mm compared to the long-term average). The annual temperature in the two experimental years was similar and approximately 0.8°C higher than the long-term average for the site (Table 1). At a monthly timescale and during part of the growing season (March to September), in comparison with the first year, the second year had a cumulated water deficit difference of −266 mm and warmer temperatures by an average of 1.9°C. Larger differences between the two years occurred in June and July with an average higher temperature (+6°C), higher water deficit (P – PET = −152.6 mm), and less precipitation (−81%) in the second year.

### Dynamics of soil temperature and relative soil water content

The soil temperature was significantly affected by the treatment, date, and treatment × date (Figure 1; Table S3). For most of the dates (February to October), the abandonment treatment corresponded with a lower soil temperature (1.76°C on average) than the cattle treatments, whereas the Ca-treatment soil temperature was significantly lower (0.64°C) than the Ca+ treatment. However, this difference was only observed to be significant for a limited number of dates in early summer of both years. The RSWC fluctuated from 0.6–0.7 at the beginning of spring to 0.38 in June during the wet year and 0.2 during the dry year, which accords with the variation of the climatic water balance (P – PET). In the case of the dry year, from summer until autumn, the RSWC remained lower than 0.4 and P – PET was negative.

### Root growth dynamics

Root growth was affected by the date and treatment × date interaction (Figure 1). Each year, the root growth peak occurred twice, in spring and autumn, and growth was markedly reduced in summer and winter. Only in the second year did growth cease in summer. Second year root growth was significantly lower than the first year. Regarding the treatment effect, the abandonment treatment showed significant lower root growth than the two cattle treatments for the spring period in both years and for the autumn period of the second year. In autumn 2014 (end of September), a delay of growth peaks was also observed, which led to a twofold higher root growth for the abandonment treatment versus the two cattle treatments. The two cattle treatments experienced similar root growth across the years and seasons.

### Seasonal root mass stock, below- and above net primary production, and root-to-shoot biomass ratio

The stock of root mass did not change throughout the seasons or across treatments (Table 2). The BNPP, ANPP, and root-to-shoot biomass ratio (R/S) were significantly lower during the second year, with a stronger effect on the BNPP (−44% on average) than the ANPP (−24% on average; Figure 2; Table 3). Only the abandonment treatment maintained its ANPP value in the second year, which led to a 48% decline in the R/S (significant treatment × year, P < 0.01; Table 3). Accordingly, a treatment effect was only observed for the BNPP during the second year, with a decline of 24% for the abandonment treatment compared to the cattle treatments. For the ANPP during the first year, the Ca+ treatment had 22% and 68% higher values, respectively, than the Ca- and abandonment treatments, while the Ca- treatment had a 38% higher ANPP than the abandonment treatment.

**Figure 2.**
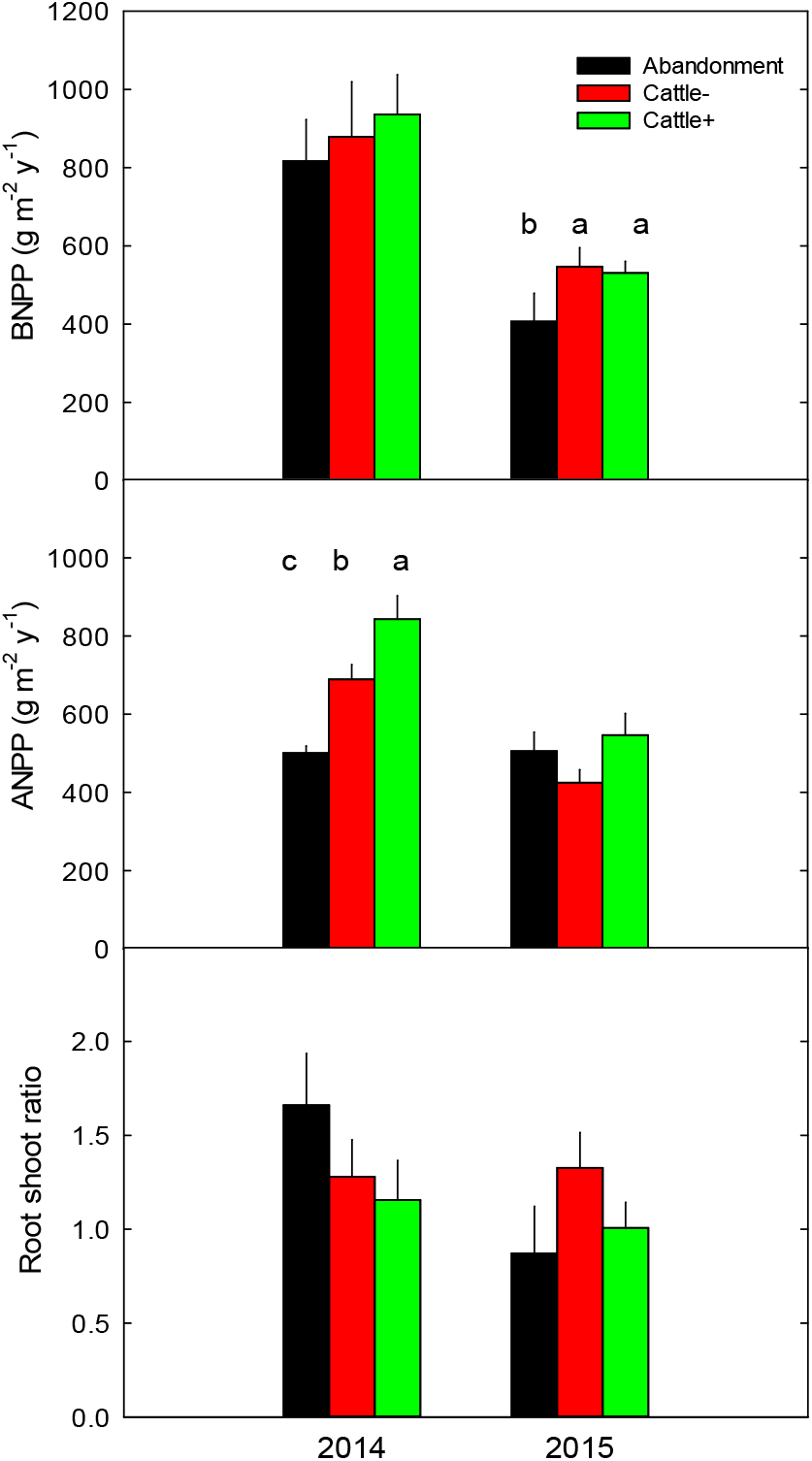
The annual root biomass production (BNPP, g m^−2^ y^−1^), annual above-ground biomass production (ANPP, g m^−2^ y^−1^), and root-to-shoot biomass ratio measured in 2014 and 2015 for the abandonment, low (Cattle-), and high (Cattle+) grazing intensity treatments. The vertical bars correspond to 1 SE (n = 4). Within a year, different letters correspond to significant differences at P < 0.05.

**Table 3.** Repeated measure ANOVA is shown for treatment, year and interaction effects on annual root production (BNPP, g m^−2^ y^−1^), annual above-ground production (ANPP, g m^−2^ y^−1^) and root to shoot ratio (R/S). Numerator (num), denominator (den) of degree of freedom (DF), *F* values are shown. Superscripts ^ns^, *,**, *** correspond to P > 0.05, P < 0.05, P < 0.01, P < 0.001, respectively.

|  |  | BNPP | ANPP | R/S |
| --- | --- | --- | --- | --- |
| | num/den DF | $F$-value | $F$-value | $F$-value |
| Treatment | 2/8 | 2.51<sup>ns</sup> | 8.10* | 0.46<sup>ns</sup> |
| Year | 1/9 | 70.72*** | 83.77*** | 13.09** |
| Treatment × Year | 2/9 | 3.83<sup>ns</sup> | 22.21** | 9.52** |

### Species composition, leaf and root traits

The abandonment treatment was characterised by the dominance of tall grass species (76% in total) with 27.2% of *Alopecurus pratensis,* 18.8% of *Elytrigia repens,* 11.3% of *Poa pratensis,* and 10.3% of *Arrhenatherum elatius.* Some forbs (19%) were also present, whereas legumes were absent (Table S2). The two cattle treatments differed from the abandonment treatment by the equal presence of *Taraxacum officinale* (18% on average) and *Trifolium repens* (17% on average). Differences also existed concerning grass species (56% in total), with the dominance of *Dactylis glomerata* (22.2%), *A. pratensis* (7.6%), and *Festuca arundinacea* (5.6%) for the Ca- treatment and *Lolium perenne* (13.6%), *D. glomerata* (9.1%), and *Poa trivialis* (7.2%) for the Ca+ treatment. Thus, the Ca+ treatment had a higher percentage of *L. perenne* than the Ca- treatment (Table S2).

The CWM leaf traits were significantly modified by the treatments. The plant height and LDMC were significantly higher (P < 0.05 and P < 0.0001, respectively) and the SLA was lower (P < 0.05) in the abandonment treatment than in the two cattle treatments (Table 4). Unlike the leaf traits, the root traits were only slightly affected by the treatments. The SRL (P < 0.1) and SRA (P < 0.05) were lower in the abandonment treatment than in the Ca- treatment, but not than in the Ca+ treatment. For other root traits (diameter, RTD, and root length percentage by class diameter), no between-treatment differences were observed (Table 4).

**Table 4.** Root traits measured from ingrowth core collected in June 2014 and leaf traits measured from botanical observation in abandonment (May 2014), Ca- and Ca+ (April 2014) treatments. Diameter: root diameter (mm); SRL: specific root length (m g^−1^); RTD: root tissue density (g cm^−3^); SRA: specific root area (m^2^ g^−1^); % 0-0.1 mm: percentage of root length in the class diameter 0-0.1 mm; % 0.1-0.2 mm: percentage of rot length in the class diameter 0.1-0.2 mm; % 0.2-0.3 mm: percentage of root length in the class diameter 0.2-0.3 mm; % > 0.3 mm: percentage of root length in the class diameter > 0.3 mm; Community-weighted mean (CWM) Height: plant height (cm); SLA: specific leaf area (cm^2^ g^−1^); LDMC: leaf dry matter content (g g^−1^). Means ± SE are shown (n = 4). num/den DF: numerator and denominator of degree of freedom. Superscripts ^ns^, ^+^, *, **, *** correspond to P > 0.1, P ≤ 0.1, P < 0.05, P < 0.01, P < 0.001, respectively. For SRL, SRA, height, SLA and LDMC, different letters correspond to significant differences between treatments.

|  | num/den<br>DF | F-value | Abandonment | Cattle- | Cattle+ |
| --- | --- | --- | --- | --- | --- |
| **Root traits** |  |  |  |  |  |
| Diameter | 2/8 | 1.61<sup>ns</sup> | 0.240 ± 0.015 | 0.210 ± 0.006 | 0.222 ± 0.015 |
| SRL | 2/8 | 3.71<sup>+</sup> | 237.2 ± 26.3 b | 332.7 ± 30.4 a | 277.8 ± 23.8 ab |
| RTD | 2/8 | 0.55<sup>ns</sup> | 0.099 ± 0.007 | 0.095 ± 0.003 | 0.102 ± 0.007 |
| SRA | 2/8 | 4.96<sup>*</sup> | 0.137 ± 0.011 b | 0.182 ± 0.008 a | 0.155 ± 0.01 ab |
| % 0-0.1 mm | 2/8 | 1.28<sup>ns</sup> | 28.5 ± 1.1 | 32.9 ± 5.5 | 28.8 ± 2.6 |
| % 0.1-0.2 mm | 2/8 | 0.46<sup>ns</sup> | 37.7 ± 4.4 | 37.7 ± 2.2 | 39.1 ± 1.8 |
| % 0.2-0.3 mm | 2/8 | 0.30<sup>ns</sup> | 16.6 ± 1.2 | 16.2 ± 2.4 | 17.1 ± 1.9 |
| % > 0.3 mm | 2/8 | 1.22<sup>ns</sup> | 17.2 ± 5.0 | 13.2 ± 1.3 | 15.1 ± 2.1 |
| **Leaf traits** |  |  |  |  |  |
| CWM_Height | 2/8 | 8.45<sup>*</sup> | 93.0 ± 3.5 a | 72.8 ± 7.0 b | 68.6 ± 3.8 b |
| CWM_SLA | 2/8 | 5.30<sup>*</sup> | 205.1 ± 5.7 b | 231.8 ± 7.3 a | 225.5 ± 7.1 ab |
| CWM_LDMC | 2/8 | 11.22<sup>***</sup> | 0.261 ± 0.008 a | 0.227 ± 0.007 b | 0.213 ± 0.010 b |

### Co-variation of traits and production

The two main axes of the standardised PCA explained 60.1% and 56.8% of the community trait and production variation in 2014 and 2015, respectively (Figure 3). For the first year, the first PCA axis (PC1), which accounted for 43.4% of the total variation, was significantly related to the leaf and root traits, ANPP, and soil temperature. The soil temperature, SRA, and ANPP had positive loadings, while the diameter, plant height, and LDMC had negative loadings (Table 5). The second PCA axis (PC2), which accounted for 16.7% of the total variation, was significantly and positively related to the root diameter and negatively related to the SRA. For the second year, the PC1 accounted for 37.4% of the total variation and was significantly related to the leaf and root traits, ANPP, and BNPP. The BNPP and SRA had negative loadings, whereas the root diameter, plant height, and ANPP had positive loadings (Table 5). The PC2, which accounted for 19.4% of the total variation, was significantly and positively related to the RSWC and the stock of root mass averaged across three dates. The abandonment treatment was significantly related to the PC1s, with negative and positive loadings for the first and second years, respectively.

**Figure 3.**
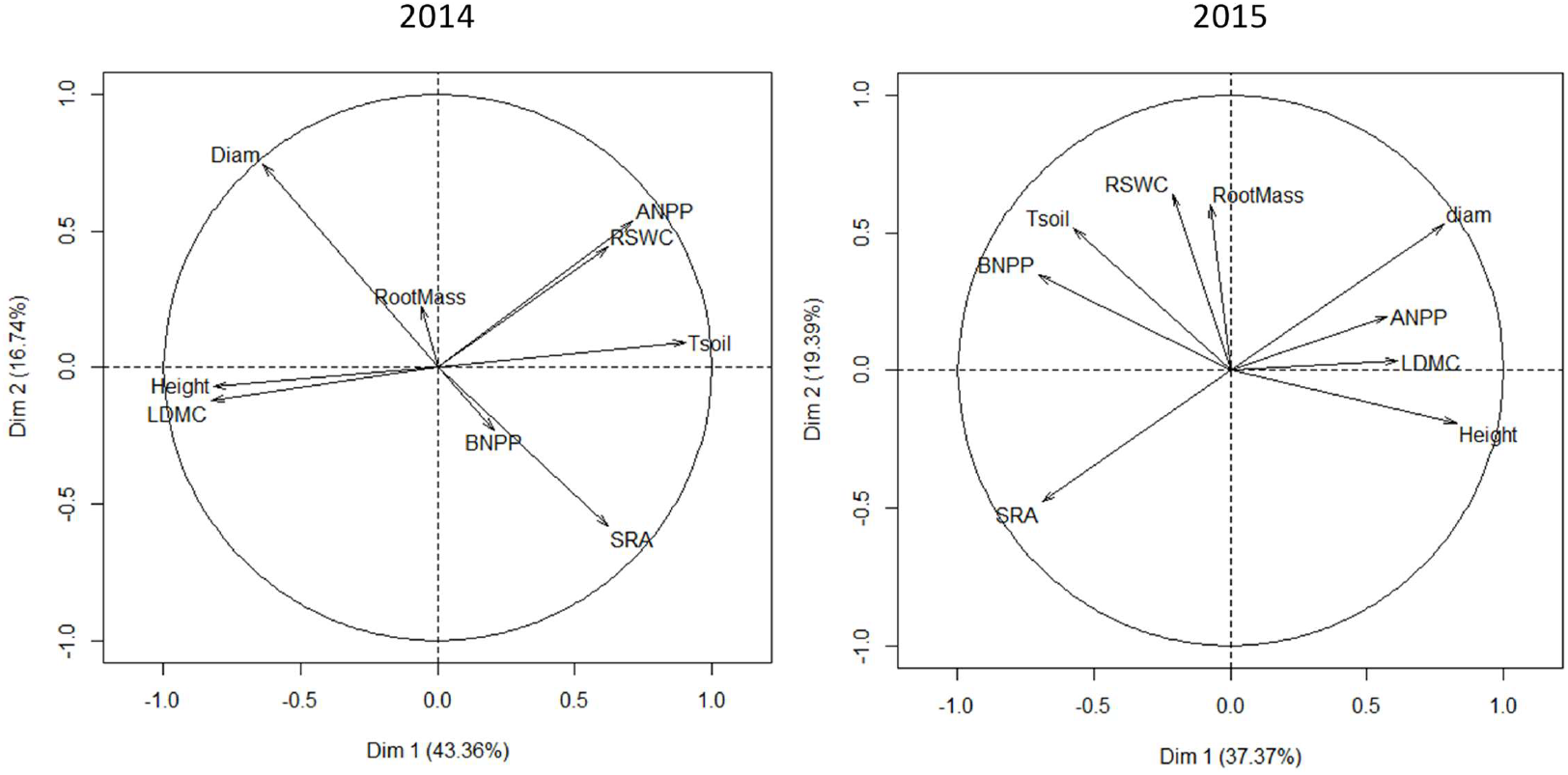
A principal component analysis (PCA) combining the leaf and root traits, above- and belowground net primary production, root mass stock, relative soil water content, and soil temperature measured in 2014 (a) and 2015 (b) for the abandonment, low (Cattle-), and high (Cattle+) stocking density treatments. Data from each plot were used in each PCA. The first two axes are shown. The arrows indicate projections of the variables within the PCA where RSWC is the relative soil water content, Tsoil is the soil temperature (°C), Diam is the root diameter (mm), SRA is the specific root area (m^2^ g^−1^), RootMass is the root mass averaged over three dates (g m^−2^), BNPP is the annual root production (g m^−2^ y^−1^), Height is the plant height (cm), LDMC is the leaf dry matter content (g g^−1^), and ANPP is the annual above-ground production (g m^−2^ y^−1^).

**Table 5.** Contribution of the different variables to the first two axes of the principal component analysis (PCA) calculated for 2014 and 2015. Variables used in the PCA were annual relative soil water content (RSWC), annual soil temperature (Tsoil, °C), root diameter (Diam, mm), specific root area (SRA, m^2^ g^−1^), root mass averaged over three dates (RootMass, g m^−2^), annual root production (BNPP, g m^−2^ y^−1^), plant height (Height, cm), leaf dry matter content (LDMC, g g^−1^), annual above-ground production (ANPP, g m^−2^ y^−1^). Treatments were added as supplementary categories. Contribution in bold indicates significant correlation of the variables on the PCA axis (P < 0.05).

| Variable | 2014 |  | 2015 |  |
| --- | --- | --- | --- | --- |
|  | Axis 1<br>(43.4 %) | Axis 2<br>(16.7 %) | Axis 1<br>(37.4 %) | Axis 2<br>(19.4 %) |
| RSWC | 0.62 | 0.44 | -0.21 | **0.64** |
| Tsoil | **0.91** | 0.09 | -0.58 | 0.52 |
| Diam | **-0.64** | **0.75** | **0.78** | 0.53 |
| SRA | **0.62** | **-0.58** | **-0.69** | -0.48 |
| RootMass | -0.06 | 0.22 | -0.07 | **0.60** |
| BNPP | 0.21 | -0.23 | **-0.71** | 0.35 |
| Height | **-0.82** | -0.07 | **0.83** | -0.19 |
| LDMC | **-0.83** | -0.12 | **0.61** | 0.03 |
| ANPP | **0.71** | 0.54 | **0.57** | 0.20 |
| *Suppl. Categories* |  |  |  |  |
| Abandonment | **-2.62** | -0.24 | **2.04** | -0.27 |
| Cattle- | 1.07 | -0.55 | -1.21 | -0.62 |
| Cattle+ | 0.70 | 0.18 | -0.83 | 0.90 |

## Discussion

Ten years of contrasted management strongly modified the functional diversity and above-ground production of this fertile upland grassland as previously shown for the same site (Herfurth et al., 2015; Louault et al., 2017). Accordingly, we expected that the above-ground biomass patterns would be mirrored below ground, especially during the periods of grazing. In this section, we first discuss the within-year differences of root growth, followed by the inter-annual variation responses to grazing intensity and climatic condition variabilities between the two contrasting years. Finally, we analyse the relationships between traits and the above- and below-ground production.

### Seasonality of root growth in relation to grazing intensity and climatic conditions

As expected, the permanent grassland root growth was affected by seasons and the spring and autumn peaks (Garcia-Pausas et al., 2011; Pilon et al., 2013; Steinaker and Wilson, 2008), but unexpectedly, grazing pressure applied by rotations and climatic condition variabilities had very limited effects on this seasonality. Accordingly, at the below-ground level, the plant community behaviour was not affected by either the rotational grazing management or the climatic condition variabilities, although a severe drought occurred in summer of the second year. Only the abandonment treatment showed a delayed root growth peak in spring. This delay was probably the result of a slower shoot budburst and a reduced capacity to produce new green leaves in the dense litter canopy, especially at the beginning of the growing season in spring (data not shown). Moreover, the tall and dense canopy of the abandonment treatment strongly modified the soil temperature; as expected, the soil conditions were cooler in the abandoned vegetation (Picon-Cochard et al., 2006; Zhou et al., 2017; Zhu et al., 2016). As shown in some studies, light, soil, water, and nutrient availability (Edwards et al., 2004; Garcia-Pausas et al., 2011; Steinaker and Wilson, 2008) are other abiotic factors determining root growth dynamics in grasslands, for example root peaks were observed before the summer soil temperature peak when a negative climatic water balance occurred, especially in the second year. Nevertheless, plants growing in the abandonment treatment offset their slower root growth by producing a similar annual root biomass, especially during the wet year. The presence of tall grass species, such as *A. pratensis, A. elatius,* and *E. repens,* having plant trait syndromes related to both disturbance and resource conservation strategies (i.e. lower SLA and SRL and higher plant height and root depth; Pagès and Picon-Cochard, 2014) might explain the tall grass capacity to produce higher root biomasses before the canopy senescence onset. Additionally, pre-existing soil fertility can be maintained in conditions of very low levels of herbage utilisation (near-abandonment) because of the absence of biomass exportation and increased internal recycling of N within senescent plants, both of which contribute to an increase in the total N available for plant growth (Loiseau et al., 2005).

The similar root growth dynamics of the two cattle treatments was unexpected since infrequent defoliation and moderate excreta returns to the soil might increase the root biomass production at the expense of the shoot biomass production (Klumpp et al., 2009). The absence of effect on the root growth and BNPP indicates that the grazing applied to the plant communities by rotations was too short but sufficient to observe the effect on the ANPP in wet conditions.

Different methods exist globally to manage grasslands by grazing (Huyghe et al., 2014), including rotational or permanent grazing options with different stocking densities, durations, and types of herbivores. In general, grazing management creates high spatial heterogeneity within the plots due to animals’ selective defoliation of plant species and because returns to soil are spatially heterogeneous. Thus, in grazed grasslands, disturbances create patches of vegetation, which should affect the local root growth and below-ground biomass of the plant communities if the intensity of grazing is sufficient. The complexity of these phenomena in grazed grasslands is greater than in mown systems (Rossignol et al., 2011).

Notwithstanding, the confounding effect of soil fertility and defoliation may mask a clear response of the below-ground compartment in grazed grasslands. In consequence, we postulate that the root growth in the Ca+ treatment was favoured by the higher soil temperature compensating for the negative effects of frequent defoliation on root growth, while the cooler soil conditions encountered in the Ca- treatment might have slowed the root growth. Soil moisture level is a main determinant of plant growth and can be affected by cattle treatments. Some studies have shown an increase of soil moisture in grazed compared ungrazed treatments due to the lower leaf area index in the grazed conditions (Moretto et al., 2001; Pineiro et al., 2010) or an absence of effect in others studies (LeCain et al., 2002; Smith et al., 2014). The presence of herbivores can increase soil bulk density and consequently modify soil moisture. In our field conditions after 10 years of treatment applications, the absence of effect on soil moisture could be due to several of these factors: absence of change of soil density, due to the rotational grazing as cattle spend less time in the plots than in continuous grazing systems; the functional composition of the community regarding both their response to defoliation and their water-use strategies; the monthly-based temporal scale used, which could buffer shorter-term responses.

We should also consider the level of soil fertility and species composition as drivers of root growth and trait plasticity (Dawson et al., 2000). The soil fertility of our site, reflected by the nitrogen nutrition index (NNI; Lemaire and Gastal, 1997), was similar across our grazing intensity gradient (Table S1), at least in 2014. Thus, our site provided us with the opportunity to compare grazing intensity effects at equivalent soil fertility levels. Because root trait plasticity generally shows larger differences with respect to soil fertility than from cutting or defoliation (Leuschner et al., 2013; Picon-Cochard et al., 2009), we could expect grazing intensity to produce a less pronounced effect on root growth under similar soil fertility conditions. Indeed, the higher presence of defoliation-tolerant species, characterised by a shorter stature and root system (e.g., *L. perenne* and *P. trivialis*) but a higher shoot and root growth capacity after defoliation and higher rhizosphere activity (Dawson et al., 2000), probably compensated for the negative effect of defoliation in the Ca+ treatment. The sampling depth might also have had an effect as we expect that due to species-specific differential root depth distribution, harvesting root systems deeper than 20 cm should have resulted in a more contrasting root growth response across the two cattle treatments according to the grass species composition (Xu et al., 2014). The results indicate that the high soil temperatures, high soil fertility, and species composition moderated the root growth response across our grazing intensity gradient. The difficulty of assigning species composition in root mixtures, however, makes it difficult to draw firm conclusions.

### Responses of ANPP, BNPP, and root-to-shoot biomass production ratio across the grazing intensity gradient and climate variability

According to meta-analyses and recent results (McSherry and Ritchie, 2013; Zeng et al., 2015; Zhou et al., 2017; Li et al., 2018), grazing intensity generally negatively affects the above- and below-ground biomass of grasslands regardless of the climatic conditions or vegetation type, although these effects can be modulated by the level of grazing intensity. The study results do not confirm these findings because the ANPP and BNPP increased in the wet and dry year, respectively, in response to grazing intensity compared to the abandonment treatment. Methodology issues for estimating the ANPP and BNPP should thus be considered for studies comparison. Some papers report the use of biomass stock or fluxes measured once at peak of growth or at several periods (Scurlock et al., 2002), other report estimation of the BNPP from indirect measurements (e.g., Zeng et al., 2015). The root mass, based on stock, provides a snapshot of plant functioning that generally includes mixtures of living and senescent tissues, thus depending on abiotic factors and plant growth, whereas measurements based on new shoot and root biomass reflect the growth potential of grasslands. Although these methods are very different, the BNPP measured with ingrowth cores produced similar results to the root mass stock assessed at three seasons. Another point to consider is the number of samples used to compare treatments and detect significant differences. In grasslands, the coefficient of variation of the root dry weight in auger samples is generally high, i.e. between 30 and 50% (Bengough et al., 2000). According to these authors, we can expect that our sampling protocol (with 16 samples) was adapted to detect at least 35% differences between treatments, whereas to detect less than 10% differences, more than 100 replicates should be collected. While it is possible that collecting more samples would have highlighted significant differences across the treatments, we had to find a compromise between performing more frequent samplings (20 dates) to study the seasonal dynamics of root growth and collecting more samples at the plot level but less frequently.

Negative climatic water balance (P – PET) conditions had stronger effects on the ANPP and BNPP than grazing intensity because severe drought had a direct negative effect on plant growth. In comparison with another experiment located along side ours, 80% of canopy senescence was reached for P – PET of −156 mm (Zwicke et al., 2013). This index reached −303 mm from March to August, confirming that a severe drought occurred in the second year of our experiment explaining the root growth cessation during the summer. At an annual scale, the ANPP of the two cattle treatments showed lower resistance to increased aridity (resistance is defined as ANPPyear2/ANPPyear1 and equalled 0.63) than the abandonment treatment (ratio = 1). For the BNPP, the results were inversed, leading to a lower resistance for the root-to-shoot biomass ratio in the abandonment treatment than in the two cattle treatments. The absence of root growth modification by grazing at an annual scale during the wet year reflects the slight change in root-to-shoot biomass allocation. Other processes, such as root turnover, are expected to change in grazed versus ungrazed grasslands. For our study, Herfurth et al. (2015) observed a similar root mass stock along a grazing disturbance gradient, but by using a simplified C flux model, these authors showed that the Ca+ treatment tended to accelerate C cycling in plant communities, resulting in a higher quantity of C allocated to the soil organic matter continuum. Taken together, these results suggest that the slight BNPP increase under grazing may occur with an increase in rhizodeposition because the root turnover, calculated as the BNPP to root mass stock ratio was not different across the treatments (data not shown; Lauenroth and Gill, 2003).

Furthermore, our results suggest that cattle treatments slow the negative effect of aridity on the root-to-shoot biomass ratio, underling that these treatments seem better adapted to buffering the negative effect of drought on grassland root production than abandonment. This finding is consistent with previous work showing that moderate grazing could be more beneficial than no grazing for drought resistance and the recovery of the ANPP and BNPP (Frank, 2007; Xu et al., 2012). The BNPP has also been shown to have greater resistant than the ANPP to changes in precipitation (Yan et al., 2013). Other studies observed no prevalent effects of grazing, drought, or fire on grassland production in North America or South Africa (Koerner and Collins, 2014). Nevertheless, further research is needed to determine whether grazing pressure has additive or combined effects on the drought response of grasslands (Ruppert et al., 2015).

### Community-weighted mean leaf and root traits as predictors of the ANPP and BNPP

As shown by some studies (e.g., Diaz et al., 2007; Laliberté and Tylianakis, 2012; Louault et al., 2017; Zheng et al., 2015), disturbances induced by grazing pressure profoundly affect plant community and functional traits by selecting defoliation-tolerant species, such as *L. perenne, P. trivialis,* or *T. repens,* with possible cascading effects on multiple ecosystem functions. Given their capacity to regrow quickly after defoliation, these species generally exhibit high values of SLA and low values of LDMC and plant height. They contrast with species that are adapted to fertile soil but exhibit a slower regrowth capacity after defoliation, such as *D. glomerata* or *F. arundinacea,* with opposite leaf trait values. In abandonment, competition for light tends to select plants with trait syndromes related to disturbance and conservative strategies (e.g., high plant height, low SLA, and high LDMC values). Thus, the CWM traits of a community depend on the balance between these species groups, which are expected to affect the ANPP and BNPP (Klumpp et al., 2009; Milchunas and Lauenroth, 1993). Although species that are tolerant and intolerant to defoliation were present in both cattle treatments, the leaf trait values were similarly and positively related to the ANPP and only differed from the traits of species present in the abandonment treatment. This means that the cessation of grazing strongly differentiated the plant communities, whereas within the two cattle treatments, differences were slighter.

For the below-ground compartment, we expected the above-ground differences to be mirrored by the root growth and traits, assuming that higher root diameter values and lower SRL and SRA values would be associated with a lower BNPP in the abandonment treatment compared with the two cattle treatments. Although the root response to grazing (mainly through defoliation) is generally reported to result in a reduction of root mass or root length (Dawson et al., 2000), our study did not confirm these assumptions. The contrasting results are possibly due to the variable abundance of defoliation-tolerant species or the combined effects of both defoliation and the level of soil fertility on the roots of grazed grasslands (Leuschner et al., 2013; Picon-Cochard et al., 2009; Yan et al., 2013; Ziter and McDougall, 2013). Thus, root growth reductions associated with grazing may have a greater impact in locations where the grazer-mediated N return is spatially decoupled from defoliation (McInenly et al., 2010). Furthermore, the higher SRA observed in the Ca- treatment than in the abandonment or Ca+ treatment should reflect a higher presence of species with fine roots, such as *D. glomerata* or *H. lanatus* (Picon-Cochard et al., 2012), because the soil fertility approximated by the NNI was similar across treatments.

### Conclusions

Rotational grazing pressures and climatic condition variabilities had limited effects on root growth seasonality, although drought had stronger effects on the BNPP than on the ANPP. The nearly identical functional traits of the plant communities and similar soil fertility across the two cattle treatments explains the absence of changes in the root mass production for these treatments. Our site disentangled the combined effects of fertility and defoliation on root production, which most previous studies have not addressed. Thus, our results suggest the prevalence of a soil fertility effect on the root production response rather than a defoliation effect. The stability of the root-to-shoot biomass ratio during the dry year evidenced a higher resistance to drought by grazed versus abandoned grassland communities. Moreover, the strong effect of climatic condition variabilities on the ANPP and BNPP observed in the short term could increase in the future as more frequent climatic extremes are expected. It is thus necessary to improve our knowledge at a larger timescale of the grazing practices that would increase grasslands resilience to more frequent and intense climatic events, such as drought and heat waves.

## Data accessibility

Data are available online: https://zenodo.org/record/4034903#.YA129-fjJPZ

## Supplementary material

**Table S1.** Nitrogen nutrition index (NNI %, Lemaire and Gastal 1997, Cruz et al., 2006) measured on forage regrowth of May in 2014 and 2015 on the non-leguminous part to assess the effect of treatments on N availability according to grazing intensity. When legumes were below 4.5% in the herbage mass, NNI was assessed using the procedure defined by Cruz et al. (2006) based on the total forage and the legume contribution. The P-values are associated with a nested mixed model: treatment used as fixed factor with plots nested in blocks as random factors. Mean ± SE is shown (n = 4). For each year, different letters correspond to significant differences at P < 0.05.

| Year | P-value | Abandonment | Cattle- | Cattle+ |
| --- | --- | --- | --- | --- |
| 2014 | 0.146 | 65.64 ± 3.10 a | 59.54 ± 1.78 a | 63.72 ± 2.86 a |
| 2015 | 0.018 | 69.72 ± 1.19 a | 61.71 ± 1.53 b | 69.25 ± 2.09 a |

---

**Table S2.** Species contribution (%) in the community present around the ingrowth core measured in April and May 2014 for Cattle-, Cattle+ and Abandonment, respectively. Mean ± SE is shown (n = 4). For each species, different letters correspond to significant differences at *: P < 0.05; **: P < 0.01; ***: P < 0.001; ns: P > 0.05.

| Group | Species | P-value | Abandonment | Cattle- | Cattle+ |
| --- | --- | --- | --- | --- | --- |
| Grasses | *Agrostis capillaris* | ns | 0.0 ± 0.0 | 0.6 ± 0.6 | 1.7 ± 1.2 |
| | *Arrhenatherum elatius* | ns | 10.3 ± 6.8 | 2.2 ± 2.2 | 2.5 ± 2.5 |
| | *Alopecurus pratensis* | ** | 27.2 ± 7.9 a | 7.8 ± 3.3 b | 3.3 ± 1.7 b |
| | *Dactylis glomerata* | * | 3.1 ± 2.7 b | 22.2 ± 9.8 a | 9.1 ± 3.8 ab |
| | *Elytrigia repens* | * | 18.8 ± 9.9 a | 2.8 ± 1.8 b | 3.8 ± 2.7 b |
| | *Festuca arundinacea*<sup>x</sup> | ns | 5.0 ± 2.3 | 5.6 ± 2.1 | 6.3 ± 2.2 |
| | *Holcus lanatus* | * | 0.0 ± 0.0 b | 4.7 ± 1.6 a | 3.4 ± 1.9 a |
| | *Lolium perenne* | *** | 0.0 ± 0.0 b | 0.9 ± 0.9 b | 13.6 ± 3.8 a |
| | *Poa pratensis* | ns | 11.3 ± 2.2 | 3.1 ± 1.5 | 3.4 ± 2.5 |
| | *Poa trivialis* | * | 0.0 ± 0.0 b | 5.0 ± 2.5 a | 7.2 ± 2.4 a |
| | *Trisetum flavescens* | ns | 0.0 ± 0.0 | 2.2 ± 1.3 | 0.6 ± 0.4 |
| Forbs | *Achillea millefolium* | ns | 1.3 ± 0.9 | 3.8 ± 2.4 | 3.1 ± 2.3 |
| | *Anthriscus sylvestris* | ns | 2.5 ± 2.1 | 0.0 ± 0.0 | 0.0 ± 0.0 |
| | *Cerastium fontanum* | ns | 0.0 ± 0.0 | 1.3 ± 0.9 | 0.0 ± 0.0 |
| | *Cerastium glomeratum* | ns | 0.0 ± 0.0 | 0.0 ± 0.0 | 0.3 ± 0.3 |
| | *Cirsium arvense* | ns | 5.0 ± 3.5 | 0.0 ± 0.0 | 0.0 ± 0.0 |
| | *Hypochoeris radicata* | ns | 0.0 ± 0.0 | 0.9 ± 0.9 | 0.0 ± 0.0 |
| | *Ranunculus acris* | ns | 0.0 ± 0.0 | 0.0 ± 0.0 | 3.8 ± 3.8 |
| | *Stellaria graminea* | ns | 0.6 ± 0.6 | 0.6 ± 0.4 | 0.0 ± 0.0 |
| | *Taraxacum officinale* agg. | ** | 0.0 ± 0.0 b | 17.5 ± 1.8 a | 19.1 ± 6.0 a |
| | *Urtica dioica* | * | 9.7 ± 4.9 a | 0.0 ± 0.0 b | 0.0 ± 0.0 b |
| | *Veronica serpyllifolia* | ns | 0.0 ± 0.0 | 0.3 ± 0.3 | 0.0 ± 0.0 |
| Legumes | *Lathyrus pratensis* | ns | 0.0 ± 0.0 | 0.3 ± 0.3 | 0.3 ± 0.3 |
| | *Trifolium pratense* | ns | 0.0 ± 0.0 | 0.0 ± 0.0 | 0.3 ± 0.3 |
| | *Trifolium repens* | *** | 0.0 ± 0.0 b | 16.3 ± 4.0 a | 17.7 ± 2.5 a |

<sup>x</sup>: new species name: *Schedonorus arundinaceus*

**Table S3.** Repeated measure ANOVA is shown for root growth (g m^−2^ day^−1^), soil temperature (Tsoil, °C) and relative soil water content (RSWC) responses to treatment, dates (d1 to d20) and interaction effects. Numerator (num), denominator (den) of degree of freedom (DF) and *F* values are shown. Superscripts ^ns^, **, *** correspond to P > 0.05, P < 0.001, P < 0.0001, respectively.

| Variables | Treatment |  | Dates |  | Treat. x Dates |  |
| --- | --- | --- | --- | --- | --- | --- |
|  | num/den DF | F-value | num/den DF | F-value | num/den DF | F-value |
| Root growth | 2/8 | 1.80<sup>ns</sup> | 19/171 | 50.40*** | 38/171 | 2.096** |
| $T_{\text{soil}}$ | 2/8 | 33.93*** | 19/166 | 944.83*** | 38/166 | 9.75*** |
| RSWC | 2/8 | 1.914<sup>ns</sup> | 19/163 | 25.287*** | 38/163 | 1.097<sup>ns</sup> |

**S4:**
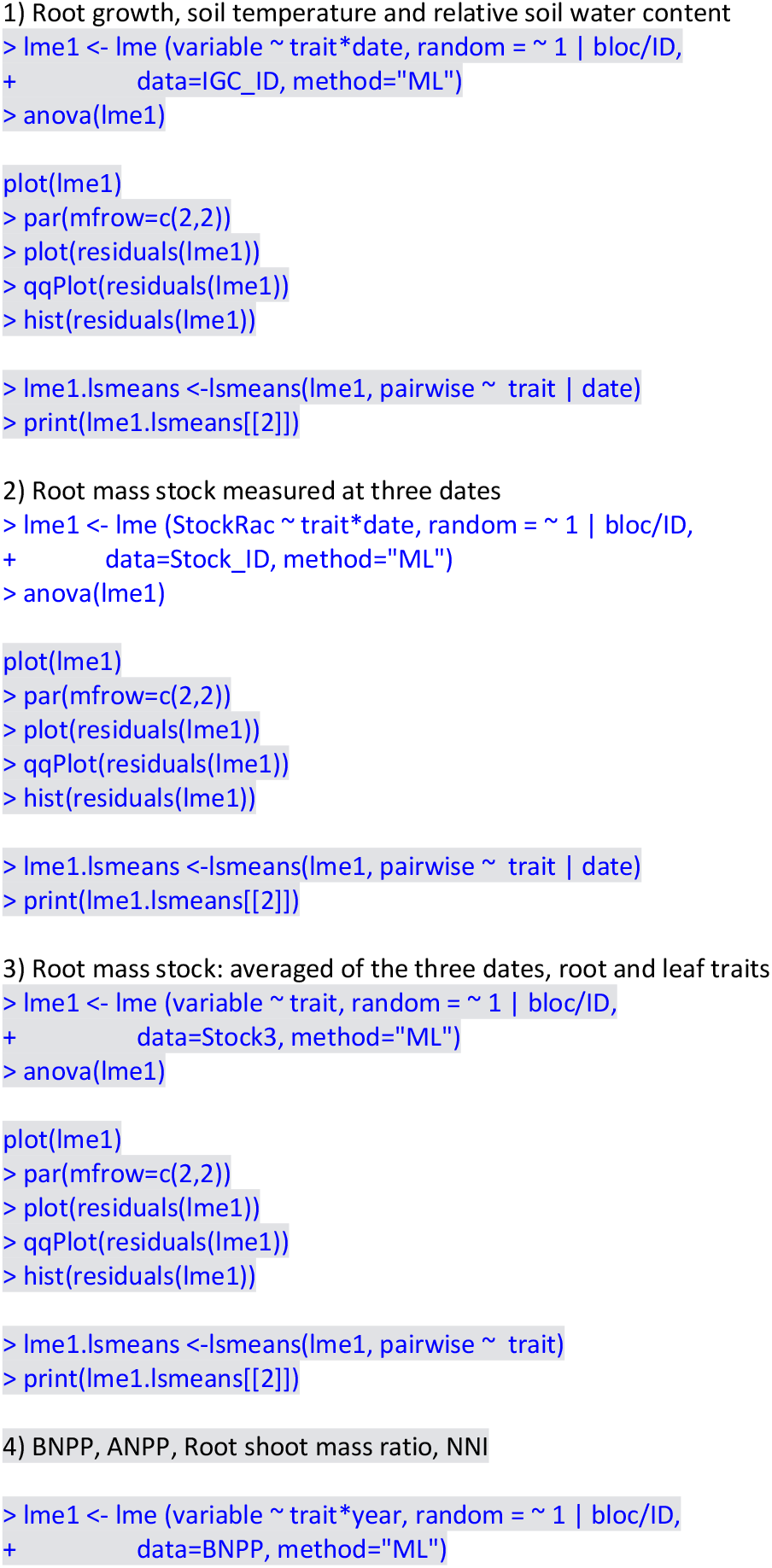
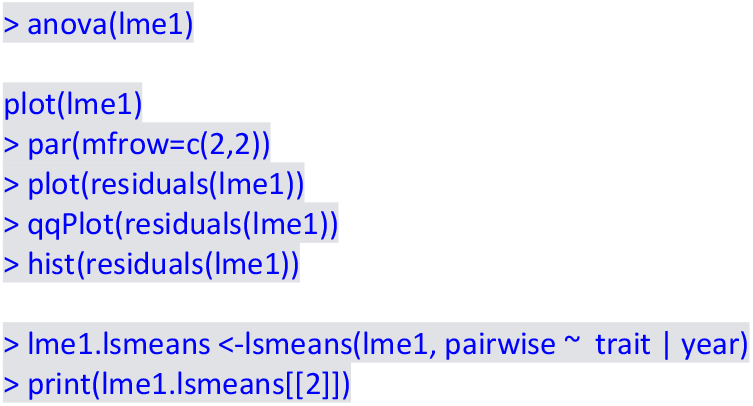
R scripts used in the paper

**Figure S1:**
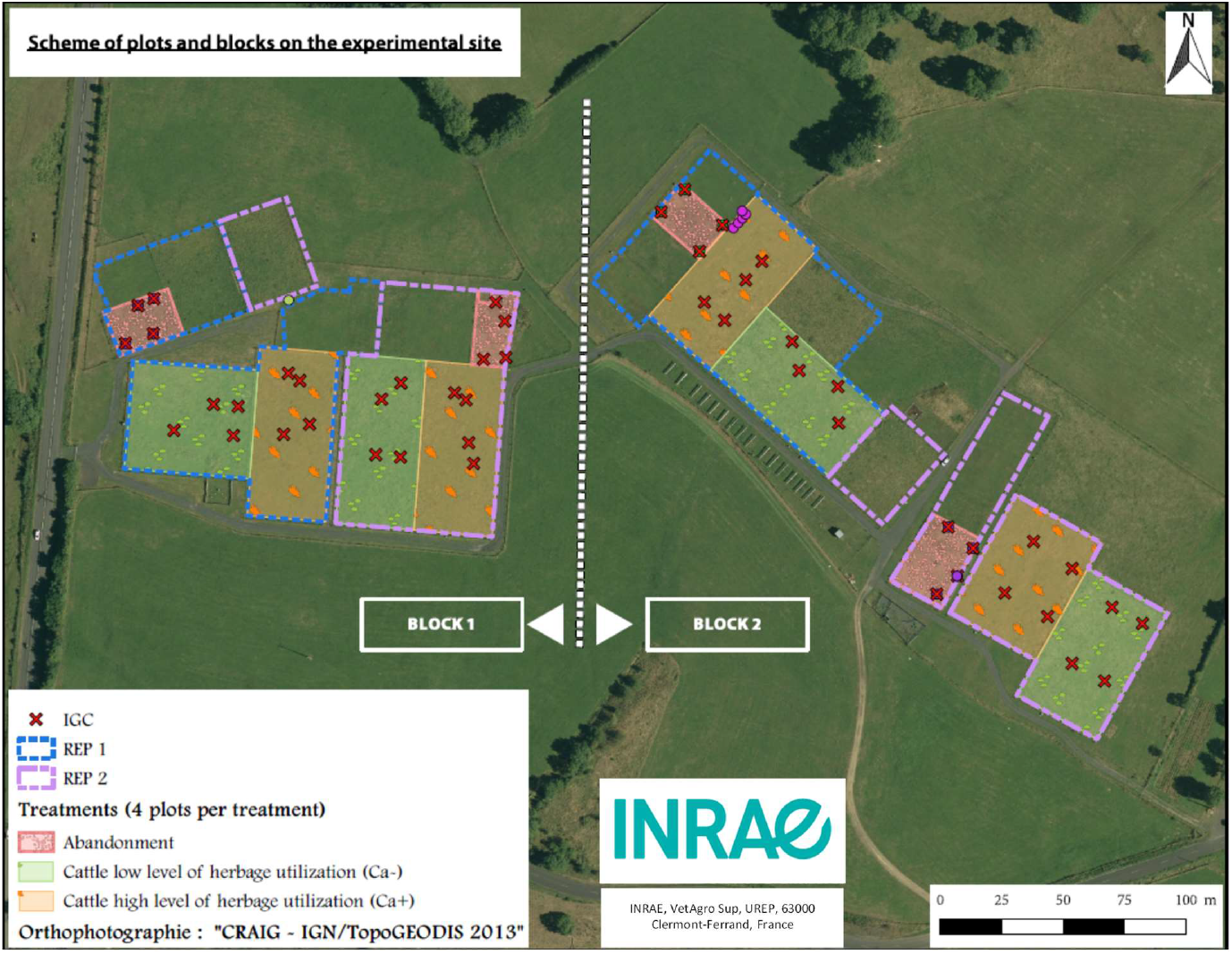
Scheme of the plots and blocks on the experimental site

## Acknowledgements

We thank staff from INRAE UREP: V. Guillot and E. Viallard for their technical expertise in field measurements, D. Colosse and S. Toillon for the soil temperature database, and S. Revaillot, A. Bartout, L. Bulon and S. Sauvat and M Mattei (VetAgro Sup) for their help in root sample measurements, and the staff of INRAE UE1414 Herbipôle. The experiment is part of the SOERE-ACBB project (http://www.soere-acbb.com/) funded by Allenvi and the French National Infrastructure AnaEE-F through ANR-11-INBS-0001. Data of the weather station are coming from the platform INRAE CLIMATIK (https://intranet.inrae.fr/climatik/, in French) managed by the AgroClim laboratory of Avignon, France. DH received a doctoral fellowship from VetAgro Sup and DGER pole “ESTIVE”. The present work falls within the thematic area of the French government IDEX-ISITE initiative 16-IDEX-0001 (CAP 20-25).

Version 6 of this preprint has been peer-reviewed and recommended by Peer Community In Ecology (https://doi.org/10.24072/pci.ecology.100073)

